# Instar determination by constrained gaussian mixture models according to Dyar’s rule

**DOI:** 10.1101/2022.12.26.521363

**Authors:** Sungmin Ji

**Affiliations:** Department of Statistics, Sungkyunkwan University, 25-2, Seonggyungwan-ro, Jongno-gu, Seoul 03063, Republic of Korea

**Keywords:** Parsimonious models, model-based clustering, delineation of developmental stage, parametric bootstraps, statistical hypothesis testing

## Abstract

Despite its importance in ecological studies and pest controls, the lack of knowledge of the life cycle and the ambiguity of data challenge the accurate determination of insect nymphs regarding many insect species. Finite mixture models are often utilized to classify instars without knowing the instar number. This study derives parsimonious gaussian mixture models using parameter constraints motivated by Dyar’s rule. Dyar’s rule explains the growth pattern of larvae and nymphs of insects by assuming a constant ratio of head capsule width for every two adjacent development stages. Accordingly, every mean value of log-transformed data in each instar stage is considered a linear function, where two Dyar constants are an intercept and a slope for the instar stages, respectively, to infer the instar stage of samples. The common variance for every instar stage regarding log-transformed data can be assumed in a mixture model, as well. If valid, these assumptions will allow an efficient estimation of the model by reducing free parameters. As a result, four model hypotheses are proposed for each assumption of instar counts depending on whether these two parameter constraints are applied. After model estimation, the proposed method uses the ICL criterion to choose the optimal counts of nymphal stages, and parametric bootstrap LR tests are applied to decide the most efficient model regarding parameter constraints. The proposed method could attain the correct model settings during the simulation study. This study also discusses the interpretation of the results of real insect data sets that concord with Dyar’s rule or not.

## 1 Introduction

Understanding the life cycle dynamics in an insect population is a significant issue in several areas of fundamental research and applied ecology, e.g., controlling the outbreak of pest insects (Logan et al. 1998). Accurate determination of the development stages of an insect species may be required in these fields. Due to an exoskeleton that limits growth, young insects grow step-wisely by their exuviation, and their life stages can be demarcated into several instar stages. However, for many species, the change in body size for successive two instar stages is often ambiguous or involves an intraspecific variation, so determining their instars is challenging. Accordingly, various clustering methods have been suggested to determine the nymphal developmental stages of an insect based on the size of morphological characters (Logan et al. 1998, Wu et al. 2013, Merville et al. 2014, Cen et al. 2018, Yang et al. 2018, Nguyen et al. 2022, Wang et al. 2022).

While some analyses for instar determination use model-free clustering methods (Yang et al. 2018, Nguyen et al. 2022), others use model-based clustering methods (Logan et al. 1998, Wu et al. 2013, Merville et al. 2014, Cen et al. 2018). One may consider a distributional assumption for each group. Finite mixture models are widely used for model-based clustering methods that assume every observation is randomly sampled from a heterogeneous population possessing several subgroups. In other words, samples can be divided into several groups, each following a different distribution. If all groups in a mixture distribution follow a normal distribution, it would be a gaussian mixture model. Let *G* be the number of groups or the number of mixture components. When it comes to univariate data, the gaussian mixture density can be written as

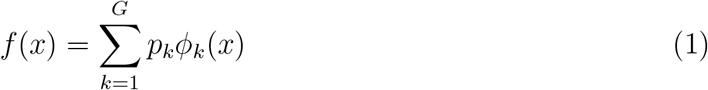

where *p*_*k*_ is the mixing proportion of *k*th group, 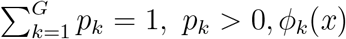 is the normal density function of 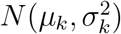. Let ***z*** = (*z*_1_, …, *z*_*G*_) be hidden group information of the sample. If ***z*** follows multinomial distribution with (1; *p*_1_, …, *p*_*G*_), i.e. *z*_*j*_ = 1 in a chance of *p*_*j*_, while *z*_*k*_ = 0 for all *k* ≠ *j*, the joint density of (*x,* ***z***) would be

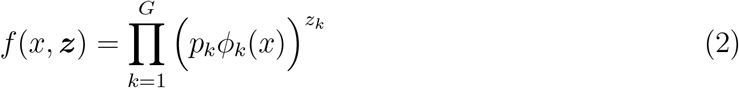

Unlike *x*, ***z*** and *G* are usually unknown and should be estimated. The expectation maximization (EM) algorithm defined in Dempster et al. (1977) is widely used to estimate ***z*** and the maximum likelihood (ML) estimate of model parameters in finite mixture models for a specified *G*. The advantage of using distributional assumptions is that it allows a solid statistical inference based on the decision theory. The mixture models can deal with classification issues in the overlapped area between two clusters with the Bayesian inference that interprets the group information of each observation as the posterior probability of belonging to a particular group given a data set (McLachlan & Peel 2000, Section 1.15). Regarding the decision theory, the highest posterior probability can be selected to determine the group for each observation. Besides, under the IID assumption—each sample is a random variable with an identical and independent mixture distribution to the other, the models allow the prediction of group information for future observations. However, the maximum likelihood estimation for gaussian mixture models often involves issues of a singularity or a spurious maximum when any pairs of sample points are sufficiently close together (Day 1969). Model constraints can deal with such issues by allowing a global maximized likelihood function (Hathaway 1985). Some parameter constraints, like common variance for every group, allow for making parsimonious models by reducing free parameters. Parsimonious models are often desirable because they provide a concise interpretation and tend to infer more precisely in model estimation and prediction when underlain assumptions are valid.

Dyar’s rule is a growth model of insects based on the assumption that indicates a regular geometric increase of head capsule width in consecutive instar stages (Dyar 1890). The model can be derived by

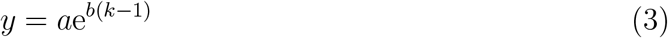

where *y* is the size of a morphological character, *a, b* are Dyar’s constants, and *k* indicates an instar stage. It implies the progression of instars will multiply e^*b*^ in the size of a morphological trait. Equation (3) is equivalent to

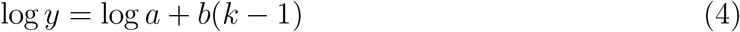

It indicates that the average size of insects in an instar can be explained by a function of Dyar constants and their developmental stage. Accordingly, conformity to Dyar’s rule can provide information on growth patterns in an insect species. Many studies about instar determination for an insect species evaluate the conformality of Dyar’s rule to their models (Logan et al. 1998, Wu et al. 2013, Sukovata 2019, Peterson et al. 2019, Nguyen et al. 2022). If the result indicates nonconformity to Dyar’s rule, it can be interpreted as underestimating or overestimating the number of instars (Sukovata 2019), or the violation of Dyar’s rule in its growth (Peterson et al. 2019).

This study suggests parsimonious gaussian mixture models for instar delineation with parameter constraints by Dyar’s rule. First, the log-transformed morphology data is assumed to follow a mixture of normal distributions. While the normality assumption of raw morphological data is frequently used in previous studies (Logan et al. 1998, Wu et al. 2013, Merville et al. 2014, Peterson et al. 2019), the log normality assumption can be realistic in practice when it comes to the length measurement because the length cannot be extended to the below zero. It often involves the right-skewed distribution whose skewness can be alleviated by log transformation. Additionally, the mean parameter of every mixture component is regarded as a linear function of Dyar’s constants and the developmental stages, just like equation (4). This assumption is likely valid when the average growth of a morphological trait in any two successive instars follows Dyar’s rule. Consequently, it allows parsimonious gaussian mixture models by replacing mean parameters with a function of two free parameters for Dyar constants. The homogeneous variance assumption for all groups also reduces free parameters. Hence, four hypotheses are suggested according to Dyar’s rule (hypothesis 1 & 2: hold; hypothesis 3 & 4: not hold) and variance homogeneity in different groups (hypothesis 1 & 3: hold; hypothesis 2 & 4: not hold) for each instar number assumption. Without the constraints over parameters, the most complex gaussian mixture model will be generated.

The gaussian mixture models can be used to determine the consistency of Dyar’s rule for an insect species with statistical hypothesis testing. That is important because while some studies suggest Dyar’s rule does not adapt to explain nymphal growth for some insect species (Peterson et al. 2019, Nguyen et al. 2022), many studies rely on a simple correlation coefficient between clustered labels and data or ratios between Dyar constants in subsequent instars (Merville et al. 2014, Sukovata 2019, Nguyen et al. 2022, Wang et al. 2022). These approaches may provide some information about instar stages but do not provide statistically powerful tests to make a decision. In contrast, the proposed method allows testing hypotheses of growth patterns in instar groups based on statistical inference if having a sufficient sample size. Suppose the test for an insect species suggests that models without Dyar’s rule outperform those resting on Dyar’s rule. It can be concluded that Dyar’s rule assumption is unreliable in modeling its instar stages in such cases. Still, such a conclusion should rely on the normality assumption of log-transformed data. Thus, in case of the violation of the distributional assumptions will be discussed at the end of the paper.

This article provides a procedure for estimating models, choosing the optimal instar counts, and testing the model hypotheses. A simulation study is presented in the result section on the performance of the proposed method. Finally, the proposed method is applied to determine the instar stages of a cicada species with a dataset collected by Hou et al. (2015).

## 2 Methods

### Constrained gaussian mixture models

Models for cluster analysis are defined as follows. From equation (1), define a mixture component density as 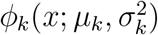 and a mixing proportion as *p*_*k*_ for *k* = 1, …, *G*. Let a vectorized mean parameter of all mixture components be ***µ*** = (***µ***_1_, …, ***µ***_*G*_)^*T*^ ∈ *R*^*G*^ and that of their variance parameters be 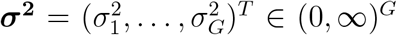. For hypotheses regarding Dyar’s rule, ***µ*** should be selected from {*M* ***β*** : ***β*** = (*β*_0_, *β*_1_)^*T*^ ∈ R^2^}, where *M* is defined as

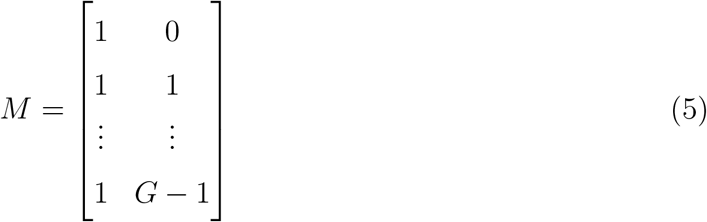

Then, ***µ*** = *M* ***β*** implies ***µ***_1_ = *β*_0_, ***µ***_2_ = *β*_0_ + *β*_1_, …, ***µ***_*G*_ = *β*_0_ + (*G* − 1)*β*_1_. For hypotheses regarding homogeneity in variance, 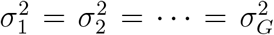, where 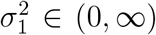. Chauveau & Hunter (2013) devised an Expectation Conditional Maximization (ECM) algorithm for gaussian mixture models with linear parameter constraints to obtain ML estimates. Accordingly, EM and ECM algorithms are proposed based on Chauveau & Hunter (2013) to obtain model estimates regarding hypotheses 1–4 for each instar number assumption. Let, *n* be the number of samples and ***x*** = (*x*_1_, …, x_*n*_). Given ***x*** with the independent sample assumption, the EM and ECM algorithms will provide 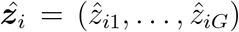, the posterior probability for hidden group information of *i*th sample given *x*_*i*_. For the clustering analysis of *x*_*i*_, the index *k ∈* {1, …, *G*} that makes the largest 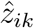 will be chosen to determine its group. The supplementary material provides details of the proposed EM and ECM algorithms.

### Instars number selection

To infer reasonable instar stages, suggested instar counts *G* are confined to a specific range while estimating models. After the estimation procedure, the optimal number of instars for each parameter constraint hypothesis will be chosen by minimizing the Integrated Classification Likelihood (ICL) derived by Biernacki et al. (2000) using the Bayesian information criterion (BIC) (Schwarz 1978). BIC and ICL are represented as equations (6) and (7).

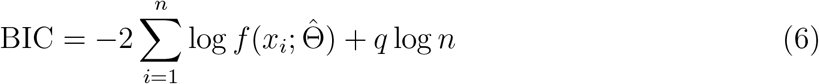

where *q* is the number of free parameters defined as

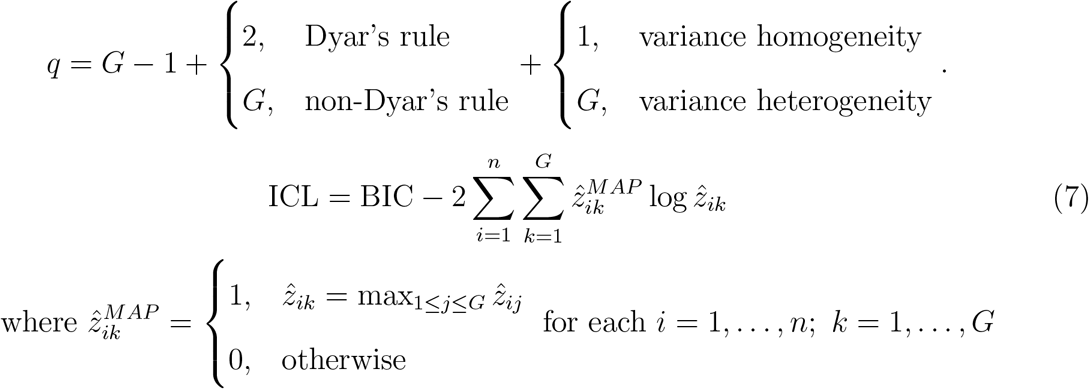

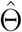 is the ML estimate of parameters given a model hypothesis, and 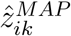 is a maximum a posteriori probability (MAP) estimate of group information for *i, k*. Suppose selected instar 7 numbers are different for each hypothesis. In that case, the final decision for the instar number can be the number held by the hypothesis with minimal ICL or in a subjective manner by interpreting the results.

### Parametric bootstrap LR tests

Statistical model assumptions enable to test of model hypotheses regarding parameter constraints. Let *ℳ*_*G*_ be a family of *G*−component gaussian mixture models regarding hypotheses 1–4, *M*_*lG*_ ∈ ℳ_*G*_ be a model for hypothesis *l, l* = 1, 2, 3, 4. To compare hypotheses 2–4 to hypothesis 1, one may consider the likelihood ratio (LR) test by calculating the maximum LR statistics given the null and alternative hypotheses. For each *M*_*iG*_ ∈ ℳ_*G*_/*M*_1*G*_

*H*_0_ : observations arise from *M*_*iG*_ versus *H*_1_ : observations arise from *M*_1*G*_

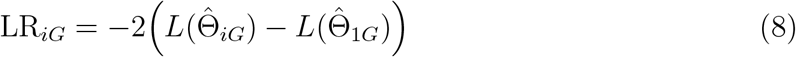

where *L*(Θ) = log *f* (***x***; Θ) is the log-likelihood function and Θ_*lG*_ are a collection of model parameters of *M*_*lG*_, for each *l* = 1, 2, 3, 4 and 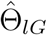 is the ML estimator of Θ_*lG*_, and LR_*iG*_ is the consequent LR statistic. The null distribution of LR_*iG*_ is usually assumed to be asymptotic *χ*^2^−distributed with *q*_LR_ = *q*_1*G*_ − *q*_*iG*_ degrees of freedom, where *q*_1*G*_ and *q*_*iG*_ denote the number of free parameters for *M*_1*G*_ and *M*_*iG*_, respectively. However, since mixture models do not guarantee the regularity condition of ML estimators, the null *χ*^2^−distribution does not provide a valid approximation for the LR statistic (McLachlan & Peel 2000, Section 6.4). As a result, conventional LR tests can lead to an inaccurate decision in comparing hypotheses, even with large sample size. A resampling method is an alternative approach to this (McLachlan 1987, McLachlan & Peel 2000, Section 6.6, Punzo et al. 2016). The parametric bootstrap LR test generates LR_*iG*_ replications as follows. *n* bootstrap samples are generated from the model *M*_*iG*_ under *H*_0_, where Θ_*iG*_ is replaced by its ML estimate calculated under *H*_0_ from the original dataset. The value of LR_*iG*_ replication is computed by fitting models *M*_*iG*_ and *M*_1*G*_ with the bootstrap samples. This process is independently repeated *R* times, and the *R* replicated values of LR_*iG*_ can approximate a null sampling distribution for LR_*iG*_ as *R* is increased. Consequently, a test can reject *H*_0_ when LR_*iG*_ for the original data overlies the specified critical region in the approximated null distribution of LR_*iG*_.

Since this process is computationally intensive when *R* is large, without an interest in estimating a precise p-value, the following criterion allows using a moderate number of *R*. Aitkin et al. (1981) note that the bootstrap replications can provide a test of approximate *α*, where *α* is a specified significance level for the test. If LR_*iG*_ for the original data is greater than the *h*th smallest of its *R* bootstrap replications, the test that rejects *H*_0_ has an approximate size of *α* as

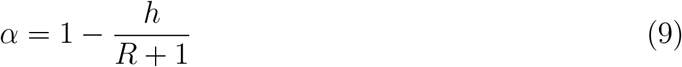

According to equation (9), for a specific *α*, the value of *R* and *h* can be chosen. For instance, when *α* = 0.01, *R* = 99 with *h* = 99 can be used to approximate *α*. Although this hypothesis testing can be applied to assess the number of *G* as well (McLachlan 1987, Hunt & Chapman 2001), this study suggests a procedure that chooses *G* with the ICL criterion and then tests hypotheses 1–4 with the parametric bootstrapping method to reduce its computation requirement. If a null hypothesis is nested into other null hypotheses, it forms a hierarchy between the null hypotheses (Figure 1). For instance, *M*_2*G*_ includes *M*_4*G*_, and this means *M*_4*G*_ is more restricted than the former. Since multiple testing can increase the Type 1 error in its decision, *α* for whole tests should be controlled by the familywise error rate. For example, the closed likelihood ratio testing procedure is applied to control the familywise error rate at level *α* (Greselin & Punzo 2013). Greselin & Punzo (2013) calculate p-values for all hypotheses in a hierarchy, like Figure 1. When they test a null hypothesis that is nested into other null hypotheses, they adjust a p-value for testing it with the maximum value of p-values regarding all LR statistics from the nested hypotheses. For example, when testing 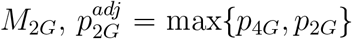 is used, where *p*_*iG*_ is a p-value of LR_*iG*_ under *H*_0_ : *M*_*iG*_. Then, *H*_0_ : *M*_4*G*_ should be tested at first when deciding on a final model because its rejection is a necessary condition for the rejection of other hypotheses. As long as *M*_4*G*_ is rejected, *H*_0_ : *M*_2*G*_ and *H*_0_ : *M*_3*G*_ are tested, otherwise *M*_4*G*_ is accepted. Two tests for *M*_2*G*_ and *M*_3*G*_ can decide the conformity to Dyar’s rule and variance homogeneity in model hypotheses. For example, if *H*_0_ : *M*_2*G*_ is rejected while *H*_0_ : *M*_3*G*_ not rejected, the final model decision is *M*_3*G*_. Table 1 represents a way to determine the final model by consecutive tests given *G*.

**Figure 1:**
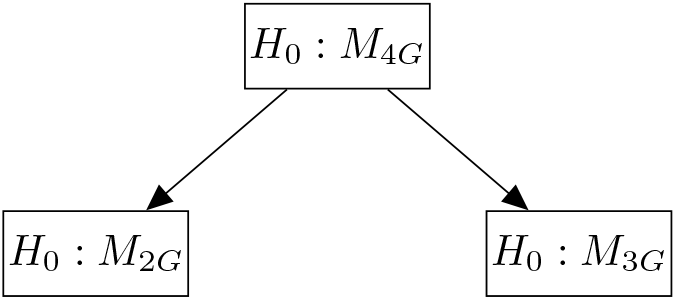
Graph of hierarchical relationships between the null hypotheses

**Table 1:** The decision for testing *H*_0_ : *M*_*iG*_, *i* = 2, 3, 4 vs. *H*_1_ : *M*_1*G*_ given an instar count *G*

| $H_0 :$ | $M_{4G}$ | $M_{2G}$ | $M_{3G}$ | | Decision |
| --- | --- | --- | --- | --- | --- |
| | NR | - | - | ⇒ | $M_{4G}$ |
| | R | NR | NR | ⇒ | $M_{4G}$ |
| | R | NR | R | ⇒ | $M_{2G}$ |
| | R | R | NR | ⇒ | $M_{3G}$ |
| | R | R | R | ⇒ | $M_{1G}$ |

NR: not rejecting $H_0$
R: rejecting $H_0$

### Simulation study

A simulation study with various situations is conducted to show the performance of the proposed procedure to find an optimal instar inference. All data analysis in this study was conducted with R programming (version 4.2.1). To assume specific cases in which a sampling distribution has an overlapped region between adjacent instar groups, the normalized overlap measure in a work of Punzo et al. (2016) is adopted. The set of overlap measures is defined by *B* = {*B*_*k,k*−1_ : *k* = 2, *…, G*}, where *B*_*k,k*−1_ is the measure of overlap between *k*th and (*k* − 1)th groups based on Bhattacharyya (1943) with a definition

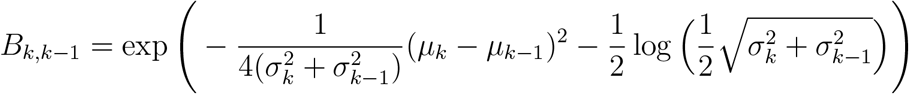

Since every *B*_*k,k*−1_ is laid between 0 and 1, max *B* = 0 implies the absence of overlapped region, whereas max *B* = 1 implies at least two adjacent groups are completely overlapped in a population. Twelve cases of *n* = 500 with various conditions regarding Dyar’s rule, variance homogeneity, overlapping between groups, and mixing proportions are considered.

For cases 1 and 3, Dyar’s rule and variance homogeneity are assumed, whereas incongruous Dyar’s rule and common variance are assumed for cases 2 and 4. Cases 1–4 are assumed to be a uniform mixture of different groups. For cases 5 and 6, groups with an unbalanced mixing proportion, Dyar’s rule, and variance homogeneity are assumed. Their mixing proportion imitates a proportion of the first to the sixth instars regarding *Blattella asahinai* data collected by Peterson et al. (2019). Cases 7–12 are variance heterogeneity versions of cases 1–6. The details of the twelve cases are noted in Table 2. One thousand datasets for each case are generated to conduct the simulation study. Figure 2 represents the theoretical distributions of twelve cases described in Table 2, where simulation data was generated.

**Figure 2:**
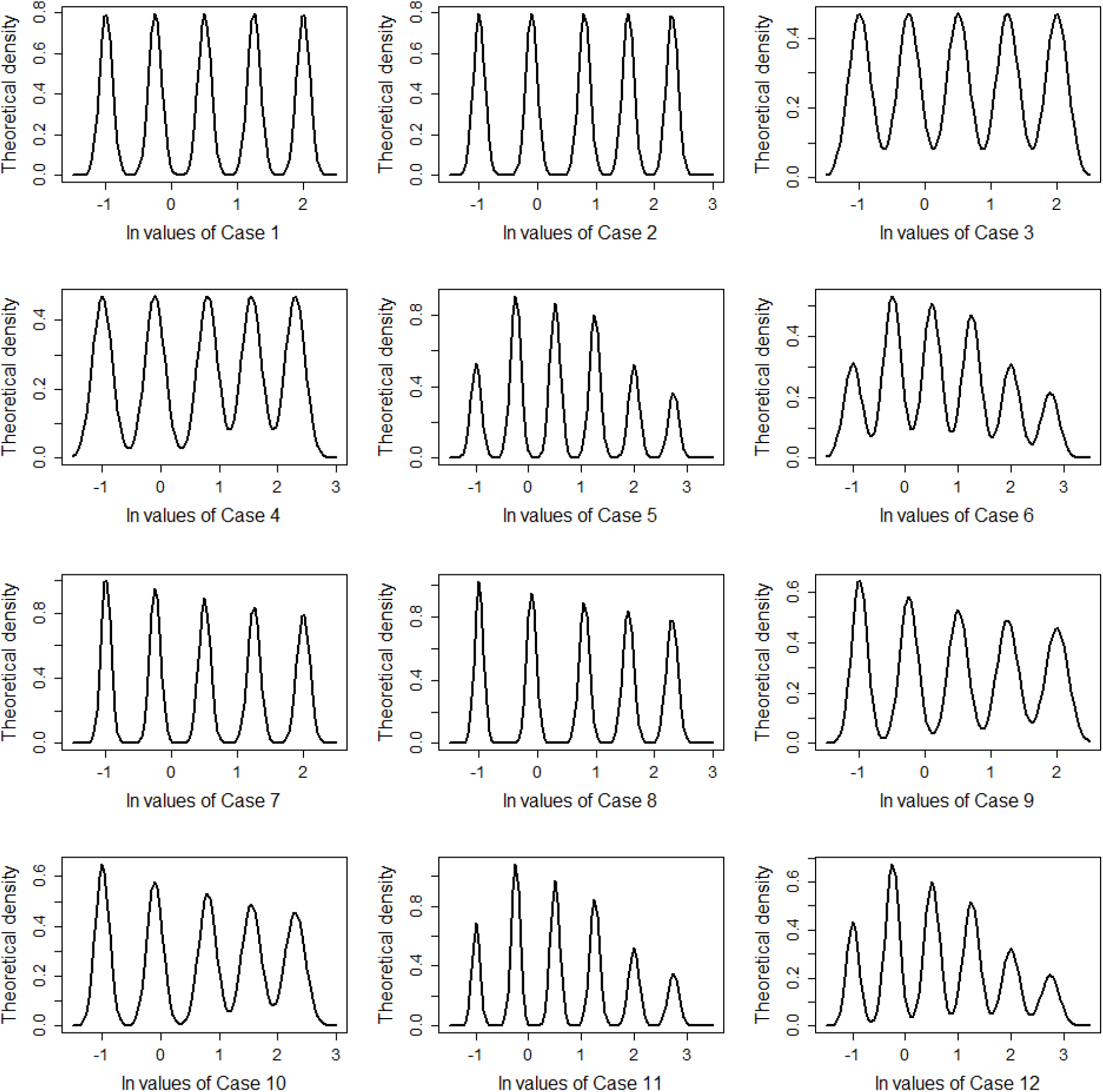
The theoretical distribution for log-transformed (ln) samples regarding cases 1–12

**Table 2:** Twelve simulation settings regarding Dyar’s rule, variance homogeneity, overlapping between groups, and mixing proportion

| No. | $G$ | $\mu^a$ | $\sigma^{2b}$ | $B^c$ | Mixing proportions<sup>d</sup> | Model |
| --- | --- | --- | --- | --- | --- | --- |
| 1 | 5 | -1/ - 0.25/0.5/1.25/2 | all 0.01 | < 0.01 | all 0.2 | $M_{4G}$ |
| 2 | | -1/ - 0.1/0.8/1.55/2.3 | all 0.01 | < 0.01 | all 0.2 | $M_{2G}$ |
| 3 | | -1/ - 0.25/0.5/1.25/2 | all 0.2875 | 0.25 | all 0.2 | $M_{4G}$ |
| 4 | | -1/ - 0.1/0.8/1.55/2.3 | all 0.2875 | 0.25 | all 0.2 | $M_{2G}$ |
| 5 | 6 | -1/ - 0.25/0.5/1.25/2/2.75 | all 0.01 | < 0.01 | 0.133/0.227/0.217/0.2/0.131/0.092 | $M_{4G}$ |
| 6 | | -1/ - 0.25/0.5/1.25/2/2.75 | all 0.02875 | 0.25 | 0.133/0.227/0.217/0.2/0.131/0.092 | $M_{4G}$ |
| 7 | 5 | -1/ - 0.25/0.5/1.25/2 | 0.006/0.007/0.008/0.009/0.01 | < 0.01 | all 0.2 | $M_{3G}$ |
| 8 | | -1/ - 0.1/0.8/1.55/2.3 | 0.006/0.007/0.008/0.009/0.01 | < 0.01 | all 0.2 | $M_{1G}$ |
| 9 | | -1/ - 0.25/0.5/1.25/2 | 0.015/0.0189/0.0227/0.0266/0.0305 | 0.05 - 0.25 | all 0.2 | $M_{3G}$ |
| 10 | | -1/ - 0.1/0.8/1.55/2.3 | 0.015/0.0189/0.0227/0.0266/0.0305 | 0.01 - 0.25 | all 0.2 | $M_{1G}$ |
| 11 | 6 | -1/ - 0.25/0.5/1.25/2/2.75 | 0.006/0.007/0.008/0.009/0.01/0.011 | < 0.01 | 0.133/0.227/0.217/0.2/0.131/0.092 | $M_{3G}$ |
| 12 | | -1/ - 0.25/0.5/1.25/2/2.75 | 0.015/0.018/0.021/0.024/0.027/0.03 | 0.05 - 0.25 | 0.133/0.227/0.217/0.2/0.131/0.092 | $M_{3G}$ |

a the mean parameter for each group
b the variance parameter for each group
c the range of $B_{k,k-1}$ in $B$
d the mixing proportion for each group

### Real data

For the application, head capsule width measurements for juveniles of insect species were collected from the literature. Hou et al. (2015) studied instar stages for *Meimuna mongolica* nymphs. They collected their samples from a laboratory and fieldwork. The first instar nymphs were raised in the laboratory, while older nymphs were excavated in the ground by fieldwork. They took pictures of the samples and measured the length of morphological characters, including head capsule and abdominal width, by Image Lab version 2.2.4.0 (resolution: 0.01 mm). Hou et al. (2015) suggested five instar stages of *M. mongolica* because according to the head capsule measurement, their samples were perfectly separated into five groups with naked eyes with no overlapping. However, this information is ignored while estimating constrained gaussian models. All the raw data were log-transformed before entering the model estimation. Due to the repeated values in datasets, the EM algorithm likely involves the singularity issue when *G* is large. Since the measurement depends on a resolution of 0.01 mm, some observations possibly have the same value, although their variable type is continuous. In dealing with such issues, raw data can be added to random noise generated with *U*_*i*_ ∼ Unif(−0.005, 0.005)mm.

## 3 Results

### Simulation study

The simulation study of the twelve cases was implemented in R. The data analysis for each case was repeated with one hundred datasets. Tables 3 and 4 represent the results of searching the optimal instar counts and model hypotheses regarding Dyar’s rule and variance homogeneity. Grey highlighted numbers imply positions of the actual model setting for each case. The ICL criterion searched for correct instar stages in 99 to 100 percent of the datasets for all cases (Table 3). However, it did not pinpoint the correct model settings for cases 7–12 that assume heterogeneous variance because it tended to select models assuming variance homogeneity. The ICL criterion is assumed to find a correct inference for instar counts, and then parametric bootstrap LR tests are conducted under the correct number of instars with 99 bootstrap replications (Table 4). With a familywise error rate as *α* = 0.01, the testing procedure selected around 98 percent of correct models assuming variance homogeneity regardless of Dyar’s rule conformity, overlapped groups, and balance in mixing proportion for cases 1–6. When it comes to variance heterogeneity data of cases 7–12, the procedure showed 15 to 63 percent to search for the correct model hypotheses. Meanwhile, regarding only being interested in Dyar’s rule consistency, the tests searched 98 to 100 percent of the correct assumption in all cases. The familywise error of 0.05 slightly increased the power of tests in cases 7–12 compared to that of 0.01 but also increased errors in Dyar’s rule conformity.

**Table 3:** Model decision of parametric LR tests given an instar count *G* for the simulation study

| Case No. | Inference of instar counts<sup>a</sup> |  |  |  |  |  | Model minimizing ICL<sup>b</sup> |  |  |  |
| --- | --- | --- | --- | --- | --- | --- | --- | --- | --- | --- |
| | 3 | 4 | 5 | 6 | 7 | 8 | $M_{1G}$ | $M_{2G}$ | $M_{3G}$ | $M_{4G}$ |
| 1 | 0 | 0 | 999<sup>c</sup> | 1 | 0 | 0 | 0 | 1 | 0 | 999<sup>d</sup> |
| 2 | 0 | 0 | 998 | 1 | 1 | 0 | 1 | 999 | 0 | 0 |
| 3 | 0 | 0 | 1000 | 0 | 0 | 0 | 0 | 1 | 0 | 999 |
| 4 | 0 | 0 | 999 | 1 | 0 | 0 | 0 | 996 | 0 | 4 |
| 5 | 0 | 0 | 0 | 1000 | 0 | 0 | 0 | 0 | 0 | 1000 |
| 6 | 0 | 0 | 0 | 1000 | 0 | 0 | 0 | 0 | 0 | 1000 |
| 7 | 0 | 0 | 998 | 1 | 0 | 1 | 1 | 1 | 34 | 964 |
| 8 | 0 | 0 | 996 | 3 | 0 | 1 | 32 | 968 | 0 | 0 |
| 9 | 0 | 0 | 994 | 6 | 0 | 0 | 0 | 0 | 125 | 875 |
| 10 | 0 | 0 | 998 | 2 | 0 | 0 | 125 | 875 | 0 | 0 |
| 11 | 0 | 0 | 0 | 1000 | 0 | 0 | 0 | 0 | 16 | 984 |
| 12 | 0 | 0 | 1 | 998 | 1 | 0 | 1 | 0 | 10 | 989 |

<sup>a</sup> instar count inference by a model that minimizes ICL in whole hypotheses
<sup>b</sup> a model minimizes ICL
<sup>c</sup> the position of grey-highlights implies the true instar number in each case
<sup>d</sup> the position of grey-highlights implies the true model in each case
Numbers in the table indicate percentages of a section for each case that refer to 1000 datasets

**Table 4:**
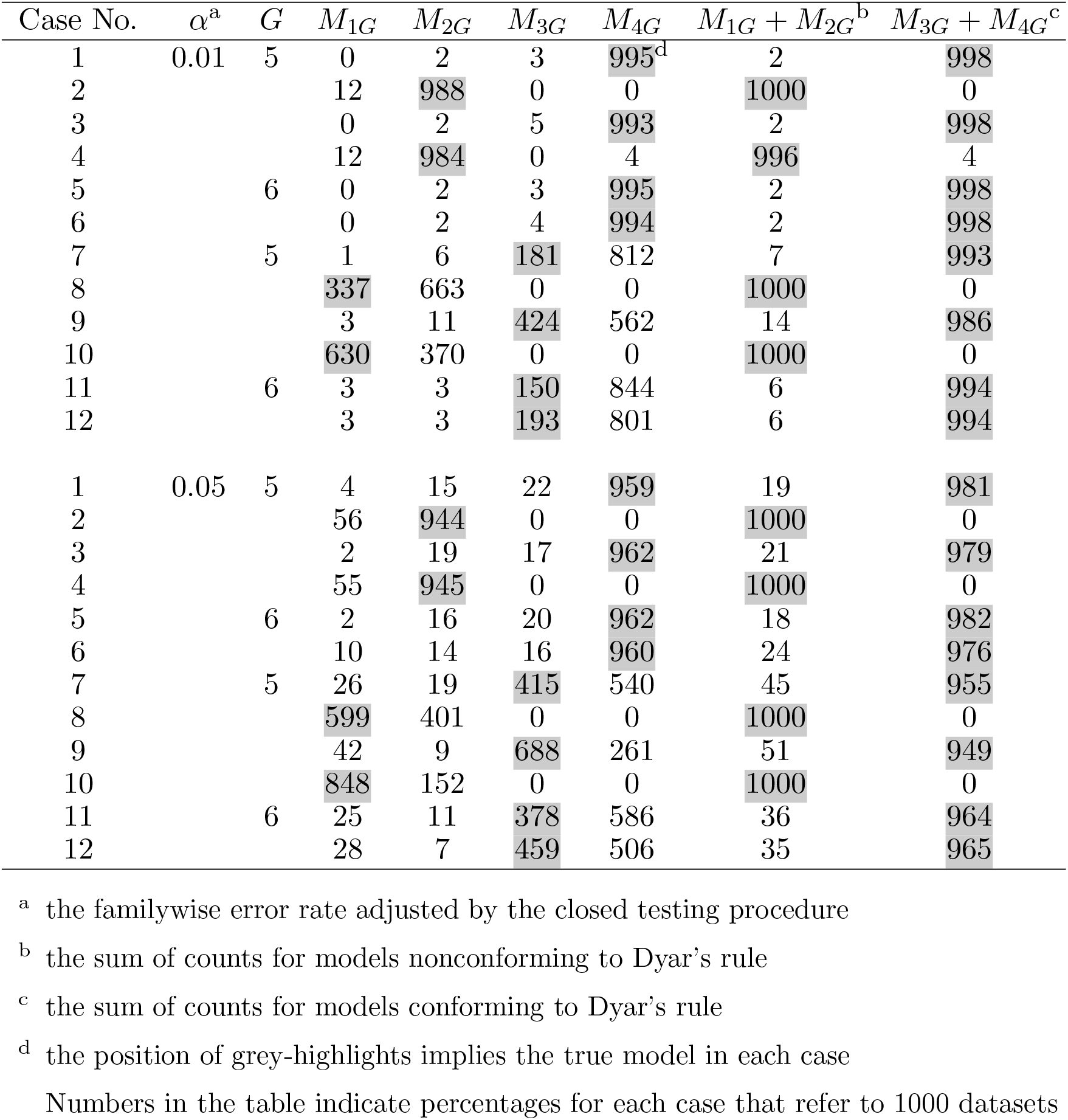
Model decision of parametric LR tests given an instar count *G* for the simulation study with 99 bootstrap replications

### Data analysis for *Meimuna mongolica*

Hypotheses 1–4 with instar counts *G* = 3, …, 8 were tested for head-capsule data of *M. mongolica*. According to the ICL criterion, the five instars assumption for *M. mongolica* was held for all the hypotheses (Figure 3). Although more parsimonious models had fewer free parameters that penalize BIC, the most complex one showed better performance in terms of ICL. Since models were well fitted with data, ICL was not significantly deviated from BIC and gave almost the same value at *G* = 5 (Figure 3 and Table 5). The estimated density of five instars for all hypotheses showed a penta-modal distribution though the second (left to right) modal had a relatively low height (Figure 4). Models that assume Dyar’s rule (*M*_3*G*_ *& M*_4*G*_) were slightly mismatched at the fourth modal of the histogram, whereas those that do not assume Dyar’s rule (*M*_1*G*_ *& M*_2*G*_) were more congruent to the histogram. The parametric bootstrap LR tests were conducted under the five-instar 17 assumption with 999 bootstrap replications. According to the test procedure, *M*_15_ was selected with a familywise error rate of 0.05 (Table 5). As a result, it can be concluded that models without assuming Dyar’s rule and homogeneous variance deliminate the instar stages of *M. mongolica* well. The clustering analysis provided the exact same result for all models assuming five instar stages.

**Figure 3:**
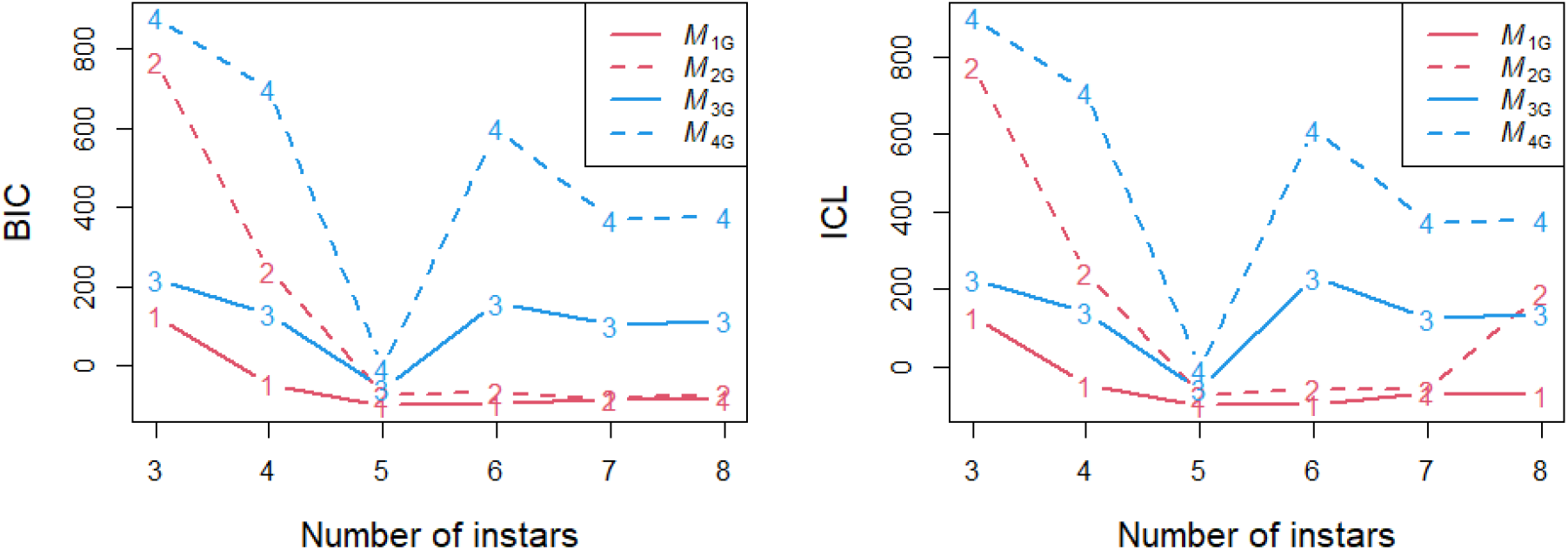
The comparison of BIC and ICL for hypothesis 1–4 of *M. mongolica* data; numbers in the plot indicate model hypothesis indices

**Figure 4:**
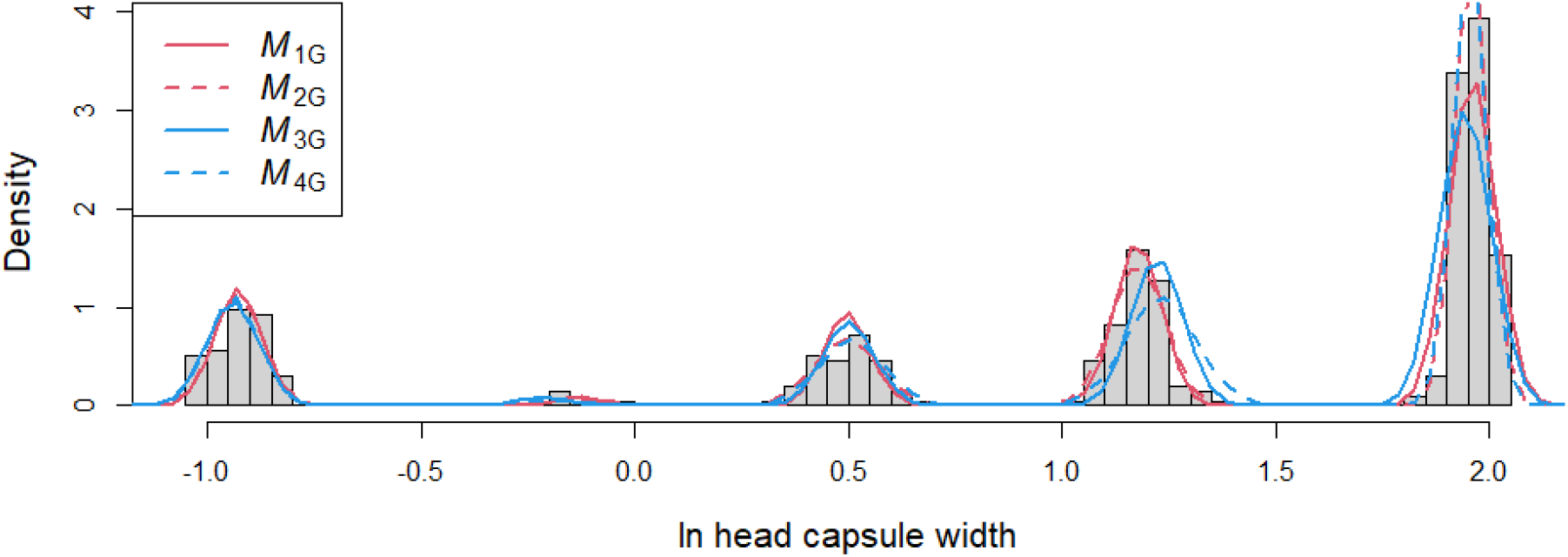
The histogram (n=391) and estimated probability density for log-transformed head capsule data of *M. mongolica* assuming five instar stages

**Table 5:** Details on the parametric LR test assuming five instar stages to the *M. mongolica* data with 999 bootstrap replications

| Model | $q^a$ | BIC | ICL | LR<sub>i5</sub> | $q_{LR}^b$ | p-value<sup>c</sup> | adj. p-value<sup>d</sup> |
| --- | --- | --- | --- | --- | --- | --- | --- |
| $M_{45}$ | 7 | -6.38 | -6.38 | 135.0 | 7 | 0.019 | 0.019 |
| $M_{35}$ | 11 | -57.92 | -57.92 | 59.5 | 3 | 0.012 | 0.019 |
| $M_{25}$ | 10 | -71.70 | -71.70 | 51.7 | 4 | 0.003 | 0.019 |
| $M_{15}$ | 14 | -99.55 | -99.55 | | | | |

a the number of free parameters for each model
b degrees of freedom of LR<sub>i5</sub> statistics
c bootstrap p-value of LR<sub>i5</sub> statistics under $M_{i5}$
d adjusted p-value by the closed testing procedure

## 4 Discussion

This study aims to make an efficient clustering procedure that determines the instar stage of insect samples by using model parsimony. Before using the proposed method, confirmation for collecting sufficient samples in every instar stage is desirable. Gaussian mixture models require sufficient samples to obtain precise estimates for their parameters. For each group of instar stages, at least two samples with different values are needed to avoid a singularity issue in calculating maximum likelihood estimates. Otherwise, the estimating procedure may not work well or lead to an erroneous outcome. Heterogeneous variances of groups can entail many local maxima in the likelihood function (Basford & McLachlan 1985) and make spurious solutions during the model estimation. Therefore, choosing an appropriate initial value for estimating models is essential to obtaining an intended result. The K-means clustering algorithm is one way to obtain a proper initial value for the ECM and EM algorithms (McLachlan & Peel 2000). The K-means initialization has been criticized for its inclination to find spherical clusters (Shireman et al. 2017), but it would not be a serious issue for univariate data, where all data clusters should be a convex set on a line. Models restricted by Dyar’s rule have an advantage in avoiding these overfitting issues compared to unconstrained models. For each group, the estimated mean parameters are hard to match a single data point because a vector for mean parameters derived from all groups should be projected in the column space of its linear constraint during the estimation process. Accordingly, they can prevent overfitting problems that fall into unintended solutions caused by a few data points. If data conforms to Dyar’s rule, they may give a more precise inference for instar stages because they use all samples to estimate the structure of a growth pattern in an insect species.

The data analysis of *M. mongolica* shows that the dataset does not match Dyar’s rule according to the parametric bootstrap tests under gaussian mixture model assumptions. Then, it shows possible discrepancies to Dyar’s rule in *M. mongolica*’s growth and gaussian mixture models without the parameter constraints seem to perform better in its instar classification. Before interpreting the result, issues involved in sampling and other issues regarding distributional assumptions should be considered. Data of Hou et al. (2015) may include heterogeneous observations. First, their observations are not the IID samples. They integrated data for the first nymphs raised in a laboratory into fieldwork data, leading to heterogeneous sampling. Besides, the fieldwork process in their study can generate a sampling bias. Except for the laboratory data, only a few small-sized cicada observations considered the second-stage instars were sampled in the fieldwork. This may indicate their methods did not adapt to collecting small-sized cicada nymphs. Therefore, more observations will be needed to evaluate Dyar’s rule in its growth and improve the accuracy of instar determination for the dataset.

While a log-normal distribution for each group is assumed in this study, violating the normality assumption can entail an inaccurate classification. Unfortunately, there is no complete method to evaluate the validity of the distributional assumption. The histogram of clustering analysis results can be used to check whether each mixture component follows a bell-shaped distribution. The log transformation possibly does not guarantee the validity of the normality assumption. Many instar determinations studies assume a normal distribution of observations for each instar stage (Logan et al. 1998, Wu et al. 2013, Merville et al. 2014), whereas the result of Peterson et al. (2019) showed that gaussian mixture models using log-transformed data provided more discrete clusters than those using untransformed data. If the log-transformation does not lead to a bell-shaped distribution for each mixture component or ambiguous clustering results, another distributional assumption can be considered for such data sets. An alternative approach for this issue is assuming a mixture density for each instar stage. Hou et al. (2015) suggested a morphological variation in *M. mongolica* that belong to the third and fourth instar stages. The difference in food availability for each place may affect their growth. The variation among habitats can affect the population’s stability, the degree of interspecific competition, and the fitness for characteristics, which may lead to morphological diversity. For some species, sexual dimorphism may contribute to the discrepancy in the data homogeneity (Turner & Williams 2005). A mixture distribution for each instar stage will represent this heterogeneity of data. Consequently, a mixture model with two or more normal mixture components instead of normal densities can be applied in the proposed procedure.

The proposed method provides an efficient way to determine instar stages under appropriate distributional assumptions. Further study is required to examine nymphal growth for various insect species and the peculiarity of the size distribution of nymphs and larvae. Accordingly, future work will investigate to relieve assumptions for adapting general cases having different distributional shapes.

## 5 Code availability

All R scripts are freely available online at the GitHub repository (https://github.com/tyrashark/cicada).

## 6 Acknowledgements

This study is conducted as an individual project in the “mixture models” lecture at Sungkyunkwan University during the fall semester of 2022. I appreciate for feedback regarding this project from Professor Byungtae Seo. I also appreciate Mr. Zehai Hou for sharing a dataset of *Meimuna mongolica* that is used to confirm the proposed procedure.

## SUPPLEMENTARY MATERIAL

### Expectation Conditional Maximization (ECM) algorithm

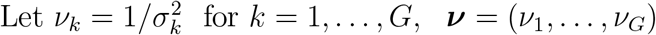

**Input: *x***, 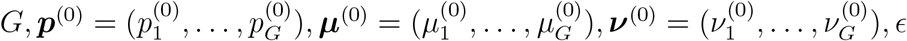

**Output** : 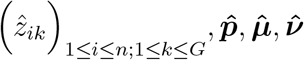

**Repeat:** for *t* = 1, …, 5000

**E-step:** Compute 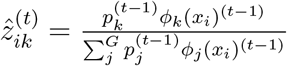 for each *i, k*

**M-step:** Update ***p***, ***µ***, ***ν*** by

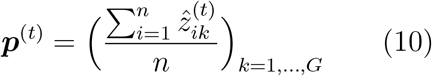

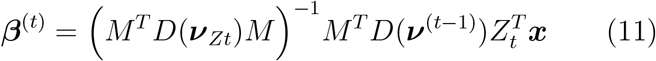

where *M* = *I*_*G*_ for hypotheses 1 & 2, equation (5) for hypotheses 3 & 4.

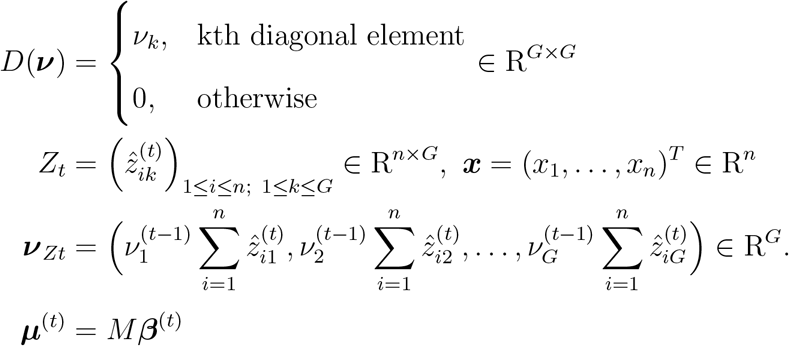

for *i* = 1, *…, n, k* = 1, *…, G*

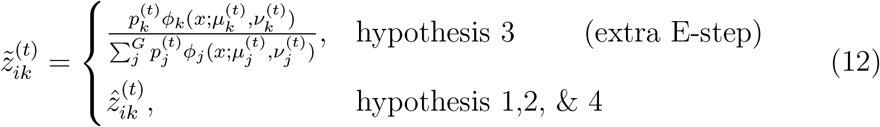

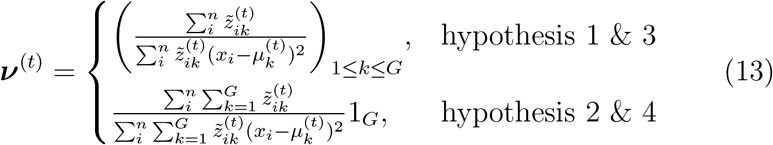

**Until:** converged in terms of *|* log *f* (***x***; ***p***^(*t*)^, ***µ***^(*t*)^, ***ν***^(*t*)^) − log *f* (***x***; ***p***^(*t*−1)^, ***µ***^(*t*−1)^, ***ν***^(*t*−1)^)*| < ϵ*

Initial value ***p***^(0)^, ***µ***^(0)^, ***ν***^(0)^ are generated by a K-means clustering algorithm that divides observations into *G* groups.

In equation (11), all 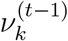 can be canceled out in *D*(***ν***_*Z*_) and *D*(***ν***^(*t*−1)^) for hypotheses 1,2, and 4, whereas 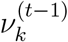 are left for hypothesis 3. This precludes obtaining a closed-form solution if updating ***p***, ***µ***, ***ν*** simultaneously during the M-step of models for hypothesis 3. Consequently, their estimation requires the ECM algorithm that fixes ***ν*** = ***ν***^(*t*−1)^ when calculating ***β***^(*t*)^ and imposes an extra E-step in equation (12) (Chauveau & Hunter 2013).

In contrast, models for other hypotheses do not need such a process, and their algorithms are equivalent to the EM algorithm.

If an algorithm is terminated due to computational issues, like an occurrence of Not-A-Number (NaN), restart the algorithm with a different initial value.

*The proof of* E-step

Under the definition of equation (1) and the IID assumption, the probability mass function of ***z***_*i*_ = (*z*_*i1*_, …, z_*iG*_) given ***x*** and Θ^(*t*−1)^ = (***p***^(*t*−1)^, ***µ***^(*t*−1)^, ***ν***^(*t*−1)^) will be

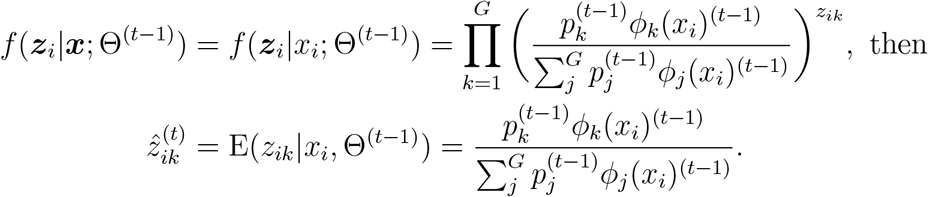

*The proof of equations* (10), (11), & (13) in M-step

Under equation (2), the complete log-likelihood is derived as

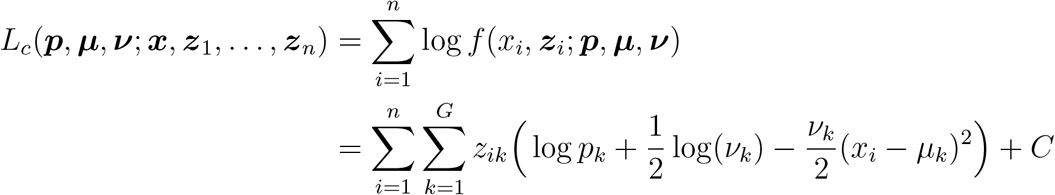

where *C* is a constant.

Given ***x*** and Θ^(*t*−1)^, the expected *L*_*c*_ is defined as

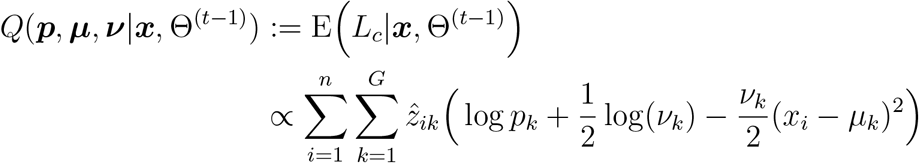

By the reparameterization of ***p*** with 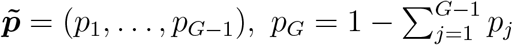

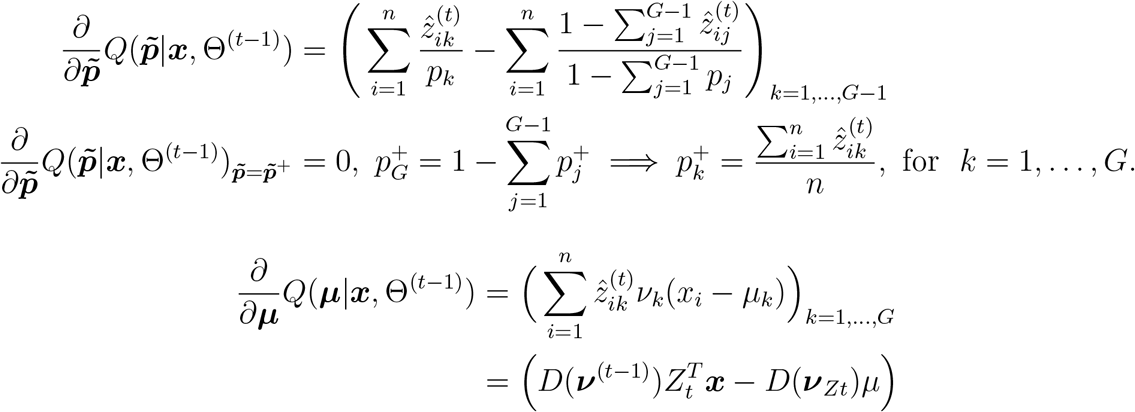

Since ***µ*** = *M* ***β***, by the chain rule

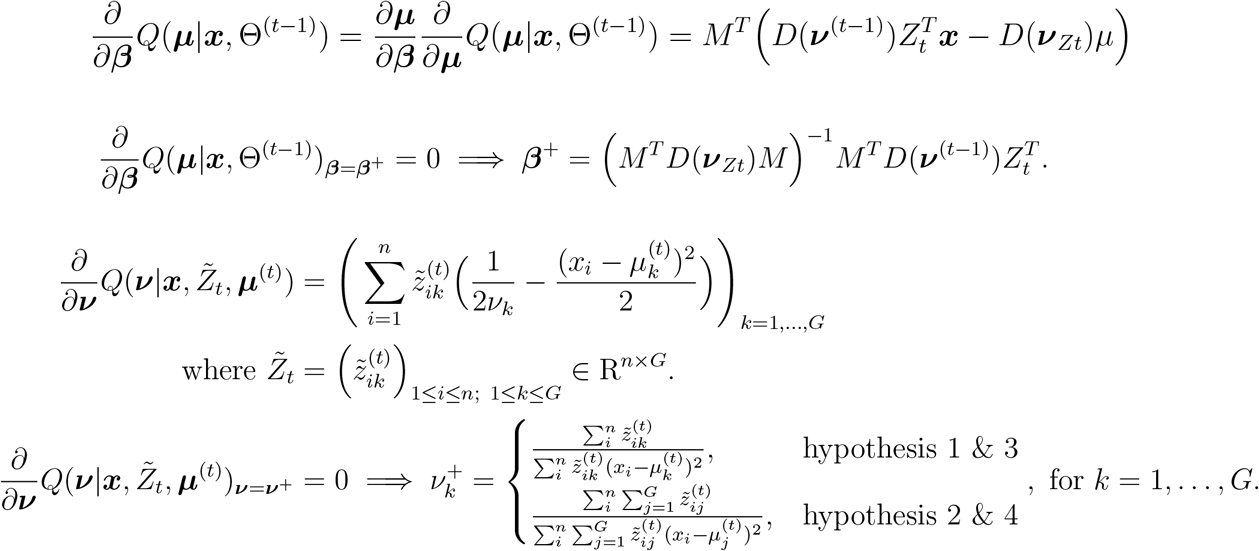

